# Sequence types of Enteroaggregative *Escherichia coli* strains recovered from human, animal, and environmental sources, India

**DOI:** 10.1101/2023.12.08.570764

**Authors:** Vinay Modgil, Harpreet Kaur, Jaspreet Mahindroo, Balvinder Mohan, Neelam Taneja

## Abstract

**Objectives:** In the current study, we report whole genome sequencing (WGS) data on EAEC strains from India to identify lineages and different sequence types (STs) in our geographical regions across North India.

**Material and methods:** We performed WGS comparative genomics characterization to examine the diversity of 122 EAEC strains collected from a large geographic area from clinical (Human sources) and non-clinical sources (animal and environmental sources). M-PCR for 21 virulence genes was performed. A triplex PCR detected phylogenetic groups A, B1, B2, and Dwas done. All strains were genome-sequenced, and bioinformatics analysis was performed.

**Results:** EAEC isolates belonged to 29 sequence types, further clustered into 11 clonal complexes, among which CC38 was the largest, containing 38 isolates mainly belonging to two ST types (ST38 and ST315). CC10 was the most diverse group, comprising 8 STs (ST43, ST2706, ST1286, ST 10, ST167, ST34, ST227, and ST4305). The most frequently detected virulence gene among the 96 clinical EAEC isolates was *ast*A (87.5%), followed by ORF3 (62.5%), and *aap* (54.1%).

**Conclusion:** These findings indicate the high diversity of EAEC and different sources of unique ST types of EAEC. Such genetic relatedness may be a favorable factor in exchanging virulence factors and other genes. The results of this study provide genetic evidence that farmed animals may act as a reservoir of EAEC.

## Introduction

In developing countries, enteroaggregative *E. coli* (EAEC) are the emerging pathogens most frequently linked to acute and chronic diarrhoea and growth retardation[1]. EAEC is also a significant contributor to severe diarrhoea in travelers from developing countries and chronic enteric infection in immunosuppressed individuals [2,3]. Various outbreaks of diarrheal disease have been attributed to it globally[4]. Notably, a significant outbreak in Germany in 2011 was caused by the hybrid O104:H4 EAEC strain, which contained the Shiga toxin gene from EHEC. This outbreak resulted in almost 4,300 cases of diarrhoea, 900 hospitalizations, and 50 fatalities [5]. EAECs are distributed widely in water and food and are occasionally found in animal feces [6].

Aggregative adherence (AA) to HEp-2 cells, a distinctive “stacked brick pattern,” was previously employed as an indicator of EAEC [7]. Adherence to the intestinal epithelium, induction of mucus production, development of biofilms, cytotoxic damage, and mucosal inflammation are all aspects of EAEC pathogenesis [8,9]. Numerous potential virulence factors have been reported [10]. Among these are the aggregative adherence fimbriae (AAF), of which five primary variations (AAF/I through AAF/V) have so far been characterized [11]. The primary transcriptional regulator AggR and the surface protein dispersin, which aid in AAF dissemination, are found on the pAA virulence plasmid, which also contains the AAFs [12]. Notably, substitute adhesins like the *E. coli* common pilus (ECP) can induce aggregative adhesion, especially when AAF is absent [10]. The cytotoxins Pet, EAEC heat-stable enterotoxin 1 (EAST-1), and hemolysinE (HlyE), which encourage epithelial damage and intestinal colonization, are additional possible virulence factors of EAEC[13][14]. Additionally, the serine protease Pic has mucinolytic properties and promotes the release of intestinal mucus [15].

Despite the discovery of several potential EAEC virulence candidates, not all EAEC strains containing these elements are harmful to humans. A single gene or set of genes has not consistently been linked by molecular tests to EAEC pathogenicity. As a result, diagnosing EAEC is still difficult and *E. coli* that lack the heat-stable or heat-labile enterotoxigenic *E. coli* (ETEC) toxins but do have the *aai*C, *agg*R, and/or *aat*A genes have been designated as EAEC.

In our geographic area, EAEC is the most prevalent diarrhoeagenic *E. coli*, with a prevalence of 11% in the pediatric age group<10 [16].EAEC was also isolated from 6% of our region’s non-diarrhoeal stools of nourished and malnourished children. Recently, Boisen *et al*. from Nataro’s group included 97 strains originating from Bangladesh, Mali, and Mozambique [17] Indian strains were not included in this study. So, in the present study, we report WGS data on EAEC strains from India to identify lineages and different STs in our regions across North India. To this aim, we have chosen a large geographical area across North India, where many laboratories have participated in this study. We employed whole genome sequencing-based comparative genomics characterization to examine the diversity of 122 EAEC strains collected from a large geographic area from clinical (Human sources) and non-clinical sources (animal and environmental sources). Whole genome sequencing (WGS) is applied to a large collection of EAECs from diverse sources for the first time. In addition, all strains were genome-sequenced, and bioinformatics analysis was used to identify their sequence types associated with disease potential. Our study provides additional information on zoonotic transmission of EAEC to humans.

## Material and Methods

### Ethical statement

The study was approved by the Postgraduate Institute of Medical Education and Research (PGIMER) Ethics Committee (INT/IEC/2017/173). Written informed consent was obtained from the patient’s parent/guardian.

### Study design

Isolates comprised of 122 strains (96 clinical and 26 non-clinical) were subjected to WGS analysis. In clinical isolates, 56 were from children with acute diarrhea aged <10 years, healthy children aged less than 5 years (n=25), malnourished children without diarrhea (n=3), immunocompromised patients with diarrhea (n=7), and chronic diarrhea (n=5). EAEC isolates of cases and controls were sourced from the previously published surveillance study from PGIMER and its 13 network laboratories in different regions across North India from 2015 to 2017. Clinical and epidemiological data was taken from the controls and the patients with diarrhea. Origin of non-clinical isolates was environmental water (n= 2), cattle stool (n=8), mutton (n=4), cheese (n=2), milk (n=3), and chutney (n=7), as shown in Table 1 of supplementary data. These samples were processed with the conventional culture method. DNA of *E. coli* isolates was extracted by heat shock method. These isolates were identified at the study site as EAEC based on PCR for the pCVD432 probe [16].

**Table 1.** Distribution of EAEC isolates among clinical and non-clinical samples used for WGS analysis.

| Source | Number of isolates |
| --- | --- |
| **Clinical EAEC isolates** |  |
| Acute diarrhea | 56 |
| Healthy Children | 25 |
| Malnourished children | 3 |
| Immunocompromised patients | 7 |
| Chronic diarrhea | 5 |
| **Non-clinical EAEC isolates** |  |
| Animal Stool and Meat | 12 |
| Food samples (Chutney, milk, and cheese) | 12 |
| Environment source (Water) | 2 |
| Total | 122 |

### Detection of virulence factors and phylogeny by M-PCR

EAEC isolates were further investigated for virulence genes. PCR for virulence genes, including *aai*C, *agg*4A, *aaf*A, *eil*A, *cap*U, *air, sat, sep*A, *sat*, ORF3, *agg*A, *ast*A, *aap, pet, sig*A, *pic, sep*A, *aaf*C, ORF61, *agg*3A, *esp*Y2, and *rmo*A was performed[18]. By amplifying the genes *chu*A, *yja*A, and a cryptic DNA fragment called TspE4C2, a triplex PCR was able to identify phylogenetic groups A, B1, B2, and D. The classification was correlated with Clermont dichotomous decision tree[16].

### Genomic DNA extraction

Complete genomic DNA was extracted from fresh overnight growth on Nutrient agar (Oxoid, Hampshire, England, U.K) using a Wizard genomic DNA purification kit (Promega Corporation, Madison, USA) according to the manufacturer’s protocol. The quality of DNA was assessed by a Nanodrop 2000 (Thermo Fisher Scientific Wilmington, DE, USA) and quantified using a Quantasfluorometer (Promega Corporation, Madison, USA). Genomic libraries were prepared using Illumina’sNextera XT DNA Library Prep kit (Illumina, San Diego, California, USA). The pooled libraries were sequenced on the IlluminaMiSeqplatform with a V3-300 reagent cartridge (Illumina, San Diego, California, USA) to generate 2x150 bp paired-end reads.

### Bioinformatic analysis for EAEC

FASTQC assessed the read quality of the genome sequence. *In silico* MLST for sequence type (ST) and clonal complex identification was performed. MLST data allele sequences and profile definitions were downloaded from the PubMLST website (pubmlst.org). Plasmid replicons and AMR gene determinants were identified using SRST2. To determine the phylogenetic relatedness between the isolates, raw reads were mapped to the reference genome of the EAEC reference strain (Accession no NC_011748) using the RedDog pipeline V1beta.10.3 (https://github.com/katholt/RedDog).Amaximumlikelihood phylogenetic tree was generated using five independent Randomized Accelerated Maximum Likelihood (RAxML) runs with a general time-reversible (GTR) substitution model and 100 bootstraps. The SNP tree with heat maps and other metadata was viewed using the Interactive Tree of Life (iTOL) online tool.

### Statistical analyses

A two-tailed chi-square test was used to compare groups. Fisher’s exact test was used if low predicted values constrained the study. The GraphPad PRISM program calculated the odds ratio (OR) and 95% confidence intervals (CIs).

## Results

### Prevalence of virulence gene and phylogroups in clinical and non-clinical isolates

The most frequently detected virulence gene among the 96 clinical EAEC isolates was *ast*A (87.5%), followed by ORF3 (62.5%), *aap* (54.1%), and *aaf*C (52%). Clinical and non-clinical isolates exhibited a high prevalence of *ast*A, with a detection rate of 87.5% and 73%, respectively. A significant difference between the virulence gene {(*agg*R (p=0.02), *aaf*C (p=0.0001), ORF61 (p=0.01), *eil*A (p=0.03), *cap*U (p=0.0005), and *rmo*A (p=0.01)} was observed between clinical EAEC isolates and non-clinical samples as shown in Table 2.

**Table 2:** Detection of virulence genes in EAEC isolates recovered from clinical and non-clinical sources.

| Virulence gene | Sampling sources (No. of isolates) |  |  |  |  |  | Sampling sources (No. of isolates) |  |  |  | P- value |
| --- | --- | --- | --- | --- | --- | --- | --- | --- | --- | --- | --- |
|  | Clinical isolates |  |  |  |  |  | Non-clinical isolates |  |  |  |  |
|  | Acute diarrhoea n=56 (%) | Healthy children (n=25) | Malnourished children (n=3) | Chronic Diarrhoea n=5 (%) | Immunocompromised patients (n=7) | Total n=96 (%) | Animal stool and meat (n=12) | Food samples (n=12) | Environment sample (n=2) | Total (n=26) |  |
| *astA* | 46 (82) | 23 (92) | 3 (100) | 5 (100) | 7 (100) | 84 (87.5) | 8 (66) | 10 (83) | 1 (50) | 19 (73) | 0.1 |
| *sigA* | 2 (3.5) | 2 (8) | 0 (0) | 1 (20) | 0 (0) | 5 (5.2) | 1 (8) | 1 (8) | 0 (0) | 2 (7.6) | 0.6 |
| *Pic* | 6 (10.7) | 3 (12) | 1 (33) | 0 (0) | 1 (14) | 11 (11.4) | 0 (0) | 1 (8) | 0 | 1 (3.8) | 0.4 |
| *sepA* | 5 (8.9) | 3 (2) | 1 (33) | 1 (20) | 1 (14) | 11 (11.4) | 1 (8) | 0 (0) | 0 | 1 (3.8) | 0.4 |
| *sat* | 10 (17.8) | 4 (16) | 1 (33) | 3 (60) | 1 (14) | 19 (19.7) | 0 (0) | 0 (0) | 0 | 0 (0) | 0.15 |
| *pet* | 5 (8.9) | 7 (28) | 1 (33) | 1 (20) | 0 (0) | 14 (14.5) | 1 (8) | 2 (16) | 0 | 3 (11.5) | 0.5 |
| **ORF 3** | 38 (67.8) | 10 (40) | 2 (66) | 5 (100) | 5 (71.4) | 60 (62.5) | 8 (66) | 3 (25) | 0 | 11 (42.3) | 0.07 |
| *aggR* | 32 (57) | 2 (8) | 2 (66) | 4 (80) | 6 (85.7) | 46 (47.9) | 0 (0) | 5 (41) | 1 (50) | 6 (23) | **0.02\*** |
| *aap* | 32 (57) | 7 (28) | 3 (100) | 5 (100) | 5 (71.4) | 52 (54.1) | 4 (33) | 6 (50) | 0 | 10 (38.4) | 0.18 |
| *aaiC* | 22 (39) | 16 (64) | 1 (33) | 1 (20) | 3 (42) | 43 (44.7) | 3 (25) | 1 (8) | 2 (100) | 6 (23) | 0.06 |
| *agg4A* | 14 (25) | 1 (4) | 2 (66) | 5 (100) | 4 (57) | 26 (27.08 ) | 0 (0) | 4 (33) | 0 | 4 (15.3) | 0.3 |
| *aggA* | 21<br>(37) | 15 (60) | 1 (33) | 0 (0) | 4 (57) | 41<br>(42.7) | 3 (25) | 4 (33) | 1 (50) | 8<br>(30.7) | 0.3 |
| *aafA* | 11<br>(19.6) | 11 (44) | 0 (0) | 0 (0) | 0 (0) | 22<br>(22.9) | 5 (41) | 1 (8) | 0 | 6 (23) | 1.0 |
| *agg3A* | 17<br>(30.3) | 8 (32) | 0 | 0 (0) | 0 (0) | 25<br>(26) | 0 (0) | 1 (8) | 1 (50) | 2 (7.6) | 0.06 |
| *aafC* | 48<br>(85.7) | 2 (8) | 0 | 0 (0) | 0 (0) | 50<br>(52) | 0 (0) | 0 (0) | 0 | 0 (0) | **0.0001\*** |
| **ORF 61** | 15<br>(26.7) | 2 (8) | 3 (100) | 5 (100) | 7 (100) | 32<br>(33.3) | 0 (0) | 2 (16) | 0 | 2 (7.6) | **0.01\*** |
| *eilA* | 27<br>(48) | 14 (56) | 1 (33) | 3 (60) | 4 (57) | 49<br>(51) | 4 (33) | 1 (8) | 0 | 5<br>(19.2) | **0.003\*** |
| *capU* | 15<br>(26.7) | 2 (8) | 2 (66) | 4 (80) | 5 (71.4) | 28<br>(29.1) | 0 (0) | 0 (0) | 0 | 0 (0) | **0.0005\*** |
| *espY* | 20<br>(35.7) | 3 (12) | 1 (33) | 4 (80) | 2 (28.5) | 30<br>(31) | 2 (16) | 1 (8) | 1 (50) | 4<br>(15.3) | 0.14 |
| *rmoA* | 24<br>(42.8) | 11 (44) | 2 (66) | 2 (40) | 3 (42) | 42<br>(43.7) | 2 (16) | 1 (8) | 1 (50) | 4<br>(15.3) | **0.01\*** |
| *shiA* | 10<br>(17.8) | 8 (32) | 0 (0) | 3 (60) | 2 (28) | 23<br>(23.9) | 2 (16) | 2 (16) | 0 | 4<br>(15.3) | 0.4 |
| *air* | 7<br>(12.5) | 7 (28) | 0 (0) | 1 (20) | 4 (57) | 19<br>(19.7) | 0 (0) | 1 (8) | 0 | 1 (3.8) | 0.07 |

\*Statistically significant ( $p < 0.005$ ) when compared pooled prevalence of EAEC virulence genes in clinical sources compared with EAEC from non-clinical sources.

Phylogenetic analysis revealed that EAEC isolates from different sources were distributed amongst the phylogenetic groups (Table 3), with subgroups D and B1 being the most prevalent among clinical isolates, containing 41.6% and 33.3% isolates, respectively. Among non-clinical isolates phylogroup B2 (42%) was most prevalent followed by B1 (19.2%). However, there was no significant difference in the distribution of phylogroup among EAEC from different sources (Table 3)

**Table 3:** Distribution of phylogenetic subgroups among EAEC isolates recovered from clinical and non-clinical sources.

| Phylogenetic group | Sampling sources (No. of isolates) |  |  |  |  | Total n=96 (%) | Sampling sources (No. of isolates) |  |  | Total n=26 (%) | P=value |
| --- | --- | --- | --- | --- | --- | --- | --- | --- | --- | --- | --- |
|  | Clinical isolates |  |  |  |  |  | Non-clinical isolates |  |  |  |  |
|  | Acute diarrhea (n=56) | Healthy children (n=25) | Malnourished children (n=3) | Chronic diarrhea n=5 (%) | Immunocompromised patients (n=7) |  | Animal stool and meat (n=12) | Food samples (n=12) | Environment sample (n=2) |  |  |
| A | 5 (8.9) | 5 (20) | 0 (0) | 0 (0) | 0 (0) | 10 (10.4) | 3 (25) | 3 (25) | 0 | 6 (23) | 0.1 |
| B1 | 18 (32) | 10 (40) | 2 (66) | 1 (20) | 1 (14) | 32 (33.3) | 1 (8) | 4 (33) | 0 | 5 (19.2) | 0.2 |
| B2 | 6 (10.7) | 3 (12) | 1 (33) | 2 (40) | 2 (28) | 14 (14.5) | 3 (12) | 1 (8) | 0 | 4 (15.3) | 1.0 |
| D | 27 (48) | 7 (28) | 0 (0) | 2 (40) | 4 (57) | 40 (41.6) | 5 (20) | 4 (33) | 2 (100) | 11 (42) | 1.0 |

\*Statistically significant ( $p < 0.005$ ) when compared pooled prevalence of EAEC phylogroups from clinical sources were compared with EAEC from non-clinical sources.

### Genetic relatedness among clinical and non-clinical isolates

Multilocus sequence (MLST) typing was used to further categories all EAEC isolates in order to establish the relationship between clinical and non-clinical EAEC strains. MLST was based on seven housekeeping genes to determine their sequence types (STs). A maximum-likelihood tree from the whole genome was constructed using EAEC prototype strain EAEC 042 as a reference (Figure 1). EAEC isolates formed tight clusters based on MLST sequence types where close clustering of EAEC isolates from the human, animal, and diverse sources was observed, as shown in Figure 1 and described in Table 4. The isolates were clustered according to their MLST STs and clonal complexes on the phylogenetic tree. Our isolates belonged to 29 sequence types, further clustered into 11 clonal complexes, among which CC38 was the largest, containing 38 isolates mainly belonging to two ST types (ST38 and ST315). CC10 was the most diverse group comprising 8 different STs (ST43, ST2706, ST1286, ST 10, ST167, ST34, ST227, and ST4305) (Table 4-6).

**Table 4.** Sequence type distribution in clinical and non-clinical/environmental EAEC isolates.

| Source | Sequence type (Number of isolates) |
| --- | --- |
| **Clinical EAEC isolates** |  |
| Acute diarrhea | **ST131**, ST200, ST328, <u>ST10(4)</u>, ST167, <u>ST34</u>, ST43 (2), ST394 (4), <u>ST130(2)</u>, <u>ST6303(2)</u>, <u>ST38 (6)</u>, <u>ST315(17)</u>, <u>ST501</u>, ST7068, <u>ST1664(2)</u> |
| Healthy Children | ST448, ST 200, ST13, ST517, **ST1286**, <u>ST10</u>, ST227, <u>ST34</u>, **ST4305**<br>ST1380, ST449, <u>ST6303(2)</u>, <u>ST38 (5)</u>, <u>ST315</u>, ST4767 |
| Malnourished children | <u>ST38</u>, ST2706, <u>ST1664</u> |
| Immunocompromised patients | <u>ST34</u>, **ST4305**, <u>ST130</u>, <u>ST6303</u>, <u>ST315</u>, <u>ST38</u>, <u>ST501</u> |
| Chronic diarrhea | **ST1588**, <u>ST501 (2)</u> |
| **Non-clinical EAEC isolates** |  |
| Animal Stool and Meat | <u>ST131</u>, ST2253, **ST1286**, <u>ST34</u>, **ST6303**, <u>ST315</u>, <u>ST38 (3)</u>, <u>ST501</u>, <u>ST1588</u> |
| Food samples (Chutney, milk, and cheese) | <u>ST131</u>, ST200, ST43, <u>ST10</u>, <u>ST38 (2)</u>, <u>ST1588</u>, ST1136, |
| Environment source (Water) | **ST4305**, ST9663 |

**Figure 1:**
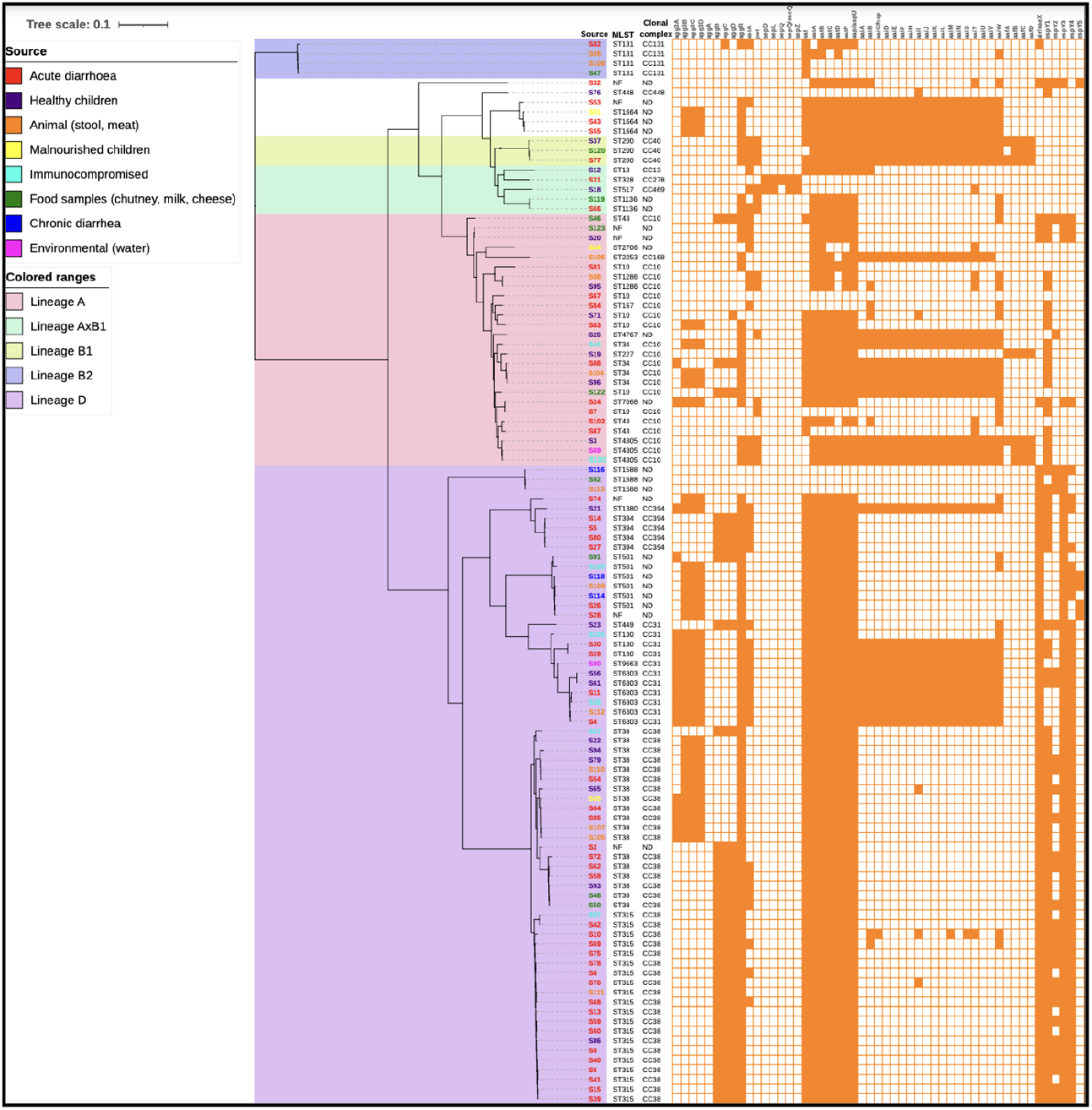
Whole-genome SNP tree for 110 EAEC isolates from various sources.

In Table 4, Sequence types that are present in both clinical and non-clinical strains are displayed in bold. For example, ST10 was found in both clinical (patients with acute diarrhoea) and non-clinical (food samples) sources. Sequence types that are shared by two or more sub-groups of the clinical or non-clinical strains are underlined. For example, ST1588 was discovered in food samples from non-clinical sources as well as in animal stools and meat.

A total of 29 different sequence types (STs) were identified. In the acute diarrheal group, a total of 15 different STs belonging to 7 different clonal complexes (CC) were identified, signifying high genotypic diversity (Table 4 and Table 5). The most common STs were ST 315 (30.3%), ST38 (10.7%), ST10 (7.14%) and ST394 (7.14%). CC38 (n=23) was the most predominant followed by CC10 (n=8) and CC394 (n=4). In 29 different sequence types identified among the 122 EAEC isolates, 11 of the isolates belonged to distinct ST type, and 15 STs contained multiple isolates, with ST315 (20 isolates), ST38 (18 isolates), and ST10 (6 isolates) being the predominant ones as shown in Table 4. The majority of EAEC isolates from non-clinical settings shared STs with clinical isolates. Importantly, our findings showed that among the non-clinical EAEC strains, 12 of the 29 STs (41%) identified in clinical isolates were also present. In addition to these, ST types ST43, ST10, ST38, and ST501 were found in food samples such chutney, milk, and cheese as well as clinical diarrheal patients. The clinical isolates ST10, ST6303, ST38, and ST315 were found in children with and without diarrhoea.In addition to the common ST types found from clinical and non-clinical EAEC, some ST types such as ST2253, ST1136, and ST9663 were detected in non-clinical sources but not among clinical sources as shown in Table 4-6.

**Table 5.** Distribution of EAEC isolates from clinical sources into sequence types and clonal complex.

| Sources | Clonal complex (CC) | Number of isolates | Sequence types (STs) |
| --- | --- | --- | --- |
| Acute diarrhea<br>N=56 | CC 131 | 1 | 131 (1) |
|  | CC40 | 1 | 200 (1) |
|  | CC278 | 1 | 328 (1) |
|  | CC10 | 8 | 10 (4), 167 (1), 34 (1), 43 (2) |
|  | CC394 | 4 | 394 (4) |
|  | CC31 | 4 | 130 (2), 6303 (2) |
|  | CC38 | 23 | 38 (6), 315 (17) |
|  | ND | 9 | 501, 7068, 1664(2) |
| Healthy Children | CC448 | 1 | 448 |
|  | CC40 | 1 | 200 |
|  | CC13 | 1 | 13 |
|  | CC469 | 1 | 517 |
|  | CC10 | 5 | 1286, 10, 227, 34, 4305 |
|  | CC394 | 1 | 1380 |
|  | CC31 | 3 | 449, 6303 (2) |
|  | CC38 | 6 | 38 (5), 315 |
|  | ND | 1 | 4767 |
| Malnourished children | CC38 | 1 | 38 |
|  | ND | 2 | 2706,1664 |
| Immunocompromised patients | CC10 | 2 | 34,4305 |
|  | CC31 | 2 | 130, 6303 |
|  | CC38 | 2 | 315, 38 |
|  | ND | 1 | 501 |
| Chronic diarrhea | ND | 3 | 1588, 501 (2) |

**Table 6.** Distribution of EAEC isolates from non-clinical sources into sequence types and clonal complex.

| Sources | Clonal complex (CC) | Number of isolates | Sequence types (STs) |
| --- | --- | --- | --- |
| Animal Stool and Meat | CC131 | 2 | 131 |
|  | CC168 | 1 | 2253 |
|  | CC10 | 2 | 1286,34 |
|  | CC31 | 1 | 6303 |
|  | CC38 | 4 | 315,38 (3) |
|  | ND | 2 | 501,1588 |
| Food samples (Chutney, milk, and cheese) | CC131 | 1 | 131 |
|  | CC40 | 1 | 200 |
|  | CC10 | 2 | 43,10 |
|  | CC38 | 2 | 38 (2) |
|  | ND | 2 | 1588,1136, |
| Environment source (Water) | CC10 | 1 | 4305 |
|  | CC31 | 1 | 9663 |

Several ST types of EAEC recovered from healthy individuals, such as ST448, ST13, ST517, ST227, ST1380, ST447, ST4767, and ST2706, were not detectable in diarrheal patients. In addition, isolates with the novel sequence types ST328, ST394, ST167, and ST7068 were isolated from acute diarrheal children, whereas an isolate with novel sequence type ST9663 was only found in the water source.

### Interpretation of the phylogenetic EAEC tree

A whole-genome SNP tree for 110 EAEC isolates from various sources was constructed using the RedDog pipeline (https://github.com/katholt/RedDog) and aligned to *E. coli* EAEC reference strain (Accession no NC_011748). A heat map of the tree with sequence types, clonal complexes, and EAEC major virulence factors was constructed on iTOL (doi: 10.1093/nar/gkz239), as shown in Figure 1. On the phylogenetic basis, our EAEC isolates are clustered into 5 lineages (A, AxB1, B1, B2, and D), further divided into multiple clades. There was intermittent clustering of isolates from different sources (clinical and non-clinical), On clade 1, four isolates of ST131, which are globally known to cause extra-intestinal infections belonging to lineage B2, formed a separate clade. These isolates were sourced from chutney samples (n=1), animals (n=2), and an acute diarrhoeal patient (n=1). On clade 2, isolates from 2 lineages, i.e., B2 and AxB1, were present. Lineage B2 in our collection was small, consisting of only three isolates, all belonging to CC40 that also carried full operon of *aat*(dispersin protein) and *aai* (secreted protein) genes. Another five isolates clustered on a branch closer to lineage B2. Lineage AxB1 consisted of five isolates from various sources (healthy, food, acute diarrhea) of four different STs. AxB1 is a hybrid phylogroup between A and B1 lineages. A single clade (clade3) consisted of isolates belonging to Lineage A (n=26) with multiple internal branching. Isolates from this clade belonged to 12 different STs, all from CC10. Isolates of ST10 were interspersed with ST7068, ST167, and ST1286, which are single-gene allelic variants of ST10, as shown in Figure 1.

Clade 4 consisted of isolates from lineage D, the largest group (N=66) in our collection. This clade further branched into three major clades (4A, 4B, and 4C). Clade 4A was formed of 3 isolates of ST1588 sourced from the animal, food samples, and a chronic diarrhea patient each. Clade 4B formed three small clusters based on the clonal complexes. Clade 4C is further comprised of two subclades where ST38 and ST315, an allelic variant of the *pur*A housekeeping gene, are divided based on sequence types. Separate branching of ST 38 and ST315 also indicates that these different STs diverged in the ancestral stage and expanded independently in our region. Isolates of ST315 were mainly from patients with acute diarrhea (17/20) and a single isolate from each healthy, immunocompromised animal. However, irrespective of the source, 75% of isolates of ST315 carried a similar virulence gene profile. Notably, isolates from chronic diarrhea patients (n=3) all belonged to lineage D.

## Discussion

The origin of pathogenic EAEC, a significant global cause of pediatric diarrhoea, is still unclear. The objective of this study was to examine the sequence types of the strains that rarely and frequently cause human diarrhoea and to estimate the relative prevalence of EAEC in clinical and non-clinical sources [19].Our findings indicated the high diversity of EAEC and EAEC from different sources of unique ST types. ST types could not be identified for 7 strains and may have new allelic variants of existing sequence types. Our study is the first to report WGS data on EAEC strains from large samples collected across North India. Recently, Boisen *et al*. from Nataro’s group included 97 strains from Bangladesh, Mali, and Mozambique [17]. Indian EAEC strains were excluded from this study. Notably, their study included clinical isolates only. The study by Nataro’s group is focused on virulence and phylogroups and not the different sequence types. In our study, 122 EAEC isolates were collected from a large geographic area, and both clinical and non-clinical settings were subjected to WGS analysis. Another major study from China reports data on 44 clinical and 33 non-clinical strains and their sequence types from Hangzhou province[20].Chattaway et al., in a previous study, analyzed 564 EAEC strains from a case-control study in Bangladesh, Nigeria, and the UK by Multi Locus Sequence Typing (MLST). They identified 17 sequence types (STs), and ST40 and ST131 were significantly associated with the disease [21].

In our findings, EAEC was highly diverse, belonging to 29 different sequence types. We found that 11 STs were less common as a single isolate of these STs was found. We found that ST315 (20 isolates), ST38 (18 isolates), and ST10 (6 isolates) were predominant. ST315 and ST38 belong to CC38, the largest in our study. A previous study from the U.K. by Anne *et al*. did not find ST38 isolate to be associated with disease or carriage [21]. This contradicts our findings, where our ST38 isolates were sourced from immunocompromised, acute diarrhea and healthy children. However, in a previous study, EAEC ST38 was associated with human ExPEC and urinary tract infections [22]. We reported that most isolates belonging to ST315 were from human cases with acute diarrhea, indicating host adaptability and high pathogenicity. We could not find any study reporting this ST type in the literature. Also, we reported that ST10 (6 isolates) and its allelic variants were present interspersed with isolates from various sources, showing it is widespread. Our results were similar to a study from China where this ST10 was detected in 5 out of 84 EAEC isolates [20]. A study from Nigeria also found ST10 to be most common and associated with a high risk of diarrhea in weaned children [23]. In previous studies, ST types, including ST10 and ST38, have been associated with EAEC disease or implicated in EAEC outbreaks [23,24]. EAEC reference strain 17–2 (from Chile) belongs to ST10, present in 6 EAEC isolates in our study. Also, EAEC ST10 was described by the pathotypes EAEC (C43/90), APEC (S17) strains, and UPEC (ATCC 23506) [25].

We have reported that transferring the EAEC from animals, food, and water sources to humans was possible. Our findings showed an overlap between clinical and non-clinical isolates; 12 out of the 29 STs (41%) detectable in clinical isolates were also detected among the non-clinical EAEC strains. In our findings, ST131 ST200, ST43, ST4305, ST1588, ST10, ST501, ST38, and S34 were detected in both clinical and non-clinical EAEC isolates, suggesting widespread reservoir/ecology and the possible transmission from environment to humans. In our findings, ST2253, ST1136, and ST9663 were detected only in non-clinical isolates, suggesting that EAEC belonging to these ST types may be more adapted to animals or environmental niches. Our study reveals that several ST types of EAEC recovered from healthy individuals, such as ST448, ST13, ST517, ST227, ST1380, ST447, ST4767, and ST2706, were not detectable in diarrheal patients, indicating that these ST types may not be pathogenic to humans. However, in the Nigerian study, ST448 was found in the diarrheal patient, indicating that there may be geographic variations, or pathogenicity may depend on host status or the presence of various virulence factors. Out of 15 STs detected in acute diarrheal patients, ST328, ST167, ST394, and ST7068 were exclusive to this group. Also, ST34, ST131, ST394, and ST38 from acute diarrheal patients were common in diarrheal patients in China [20]. However, these STs were not identified in the Nigerian EAEC isolates. This may be due to geographical restrictions in Asia [26]. ST2253 detected from an animal source (meat) in our study was also recovered from healthy humans from China, which indicates the widespread nature of this ST type and potential zoonotic source of infection [20].

In our findings, the sequence type of ST38 and ST501, which were found to cause acute and chronic diarrhea, was also recoverable from non-clinical food samples. Likewise, the sequence types of ST131 and ST1588 were detected in acute diarrheal patients, animal sources, and food samples (chutney), suggesting transmission from animal /environment due to the unhygienic preparation of the food. ST131, ST34, ST6303, ST315, ST38, and ST501 were detected in clinical acute diarrheal patients and animal and meat samples, suggesting the possible transmission route from animals. Several ST types of EAEC recovered from healthy individuals, such as ST448, ST13, ST517, ST227, ST1380, ST447, ST4767, and ST2706, were not detectable in diarrheal patients, indicating that these ST types may not be pathogenic to humans. The possibility that bacteria from the faeces were transported into the adjacent rivers and then created a much broader region of pollution may help to explain this occurrence. Overlap of identical STs in clinical and non-clinical samples shows a high probability that human infections arose from these non-clinical sources.

Our is the first study from India to report ST131 in non-bloodstream infection. Globally, ST131 is known as the multidrug-resistant (MDR) extraintestinal pathogenic *E. coli* (ExPEC) lineage, causing a large spectrum of diseases, like infections of the blood and (urinary tract infections) UTIs[27,28]. Interestingly, ST131 was not detected in healthy (nourished and malnourished children), whereas it was present in acute diarrheal, animal, and food sources, suggesting its pathogenicity and the possible transmission of this clinically important ST type. Although cattle feces are known to be the most important source of pathogenic *E. coli* in dairy farms that could contaminate raw milk, there are controversial data on animal reservoirs of EAEC. Little data shows the absence of the *agg*R gene in animal strains. Our study found a mixed proportion of animal isolates with the presence or absence of the *agg*R gene. All meat samples were positive for *agg*R, indicating that either the meat was contaminated from human sources or even animal stools, as some of the animal stool isolates also had *agg*R. The majority of EAEC isolates from phylogroups A and B1 are thought to be commensal strains, whereas subgroup B2 is most commonly linked to extra intestinal infections.

Similar to a research conducted in China, we did not discover a significant variation in the distribution of phylogenetic groups of EAEC between clinical and non-clinical strains. The study’s interesting finding is the similar percentage of B2 EAEC isolates from clinical isolates (14.5%) and non-clinical isolates (15.3%). Nevertheless, the concept of unequal virulence between clinical and non-clinical EAEC strains is compatible with our finding of a discrepancy in the patterns of distribution of known virulence genes across clinical and environmental isolates.

Among the clinical and nonclinical EAEC isolates, the *ast*A gene, which encodes the EAST-1 toxin, was most predominant and was detected in 87.5% and 73% of the isolates, respectively. This finding agrees with numerous other studies, which showed that isolates recovered from clinical and non-clinical animal samples frequently carried this virulence gene [29–31]. Our finding that *ast*A is widely distributed among *E. coli* isolates recovered from both sources suggests that all *E. coli* strains can theoretically cause diarrhoea, but the severity of the illness may depend on the gene copy number, expression level, and degree of coexistence with other virulence elements. The *ast*A gene alone has been reported to cause diarrhoea in Japan. It should be highlighted that virulence genes have been identified substantially more frequently in organisms recovered from clinical sources than in non-clinical isolates. Our findings showed that other virulence factors, including *aap,aat*A, and *agg*R, may be the distinguishing characteristics of clinical EAEC that causes human diarrhoea.

The findings show that EAEC has a high degree of variety and that its distinctive ST types are derived from several sources. According to our findings, a variety of STs contribute to the diversity of EAEC, and comparable STs in clinical and non-clinical samples had relatively close evolutionary distances. The exchange of virulence factors and other genes may be facilitated by such genetic relatedness.In summary, this work uses MLST analysis to offer an overview of the prevalence and genetic diversity of EAEC from human and other sources. The findings of this study offer genetic evidence that domesticated animals may serve as a reservoir for EAEC. Our findings add to the body of knowledge by demonstrating that animals are probably an EAEC reservoir. According to the study’s results, EAEC strains were found in both non-clinical and clinical settings in a variety of ways, and some clinical isolates may have come from non-clinical settings.

## Declarations

### Funding

None

## Data availability

The authors confirm that all data, code, and protocols have been provided within the article or through supplementary data files.

## Author Contributions

VM, Writing – original draft, VM, and NT, Conceptualization.VM, Data curation.VM and HK, Formal analysis. VM and HK, Investigation. VM and HK, Methodology. VM,JM and HK, Software. BM and NT, Supervision. VM, HK, and NT, Validation. VM, HK, and NT Writing – review and editing.

## Consent to participate

Not applicable.

## Consent for publication

Not applicable.

## Competing interests

The authors declare no competing interests.

## Acknowledgment

We thank the Director of PGIMER Chandigarh for making this effort possible. The first author acknowledges the assistance of the UGC, New Delhi, which is provided as a fellowship (Sr. No. 20614305077, Ref No. 22/06/2014(i)EU-V).

## Notes

### Competing Interest Statement

Conflict of Interest: None

